# Expansion of *in vitro Toxoplasma gondii* cysts using enzymatically enhanced ultrastructure expansion microscopy

**DOI:** 10.1101/2024.04.24.590991

**Authors:** Kseniia Bondarenko, Floriane Limoge, Kayvon Pedram, Mathieu Gissot, Joanna C. Young

## Abstract

Expansion microscopy (ExM) is an innovative approach to achieve super-resolution images without using super-resolution microscopes, based on the physical expansion of the sample. The advent of ExM has unlocked super-resolution imaging for a broader scientific circle, lowering the cost and entry skill requirements to the field. One of its branches, ultrastructure ExM (U-ExM), has become popular among research groups studying Apicomplexan parasites, including the acute stage of *Toxoplasma gondii* infection. The chronic cyst-forming stage of *Toxoplasma*, however, resists U-ExM expansion, impeding precise protein localisation. Here, we solve the *in vitro* cyst’s resistance to denaturation required for successful U-ExM of the encapsulated parasites. As the cyst’s main structural protein CST1 contains a mucin domain, we added an enzymatic digestion step using the pan-mucinase StcE prior to the expansion protocol. This allowed full expansion of the cysts in fibroblasts and primary neuronal cell culture without interference with the epitopes of the cyst-wall associated proteins. Using StcE-enhanced U-ExM, we clarified the shape and location of the GRA2 protein important for establishing a normal cyst. Expanded cysts revealed GRA2 granules spanning across the cyst wall, with a notable presence observed outside on both sides of the CST1-positive layer.

**Importance:** *Toxoplasma gondii* is an intracellular parasite capable of establishing long-term chronic infection in nearly all warm-blooded animals. During the chronic stage, parasites encapsulate into cysts in a wide range of tissues but particularly in neurons of the central nervous system and in skeletal muscle. Current anti-Toxoplasma drugs do not eradicate chronic parasites and leave behind a reservoir of infection. As the cyst is critical for both transmission and pathology of the disease, we need to understand more fully the biology of the cyst and its vulnerabilities.

The advent of a new super-resolution approach called ultrastructure expansion microscopy allowed in-depth studies of the acute stage of *Toxoplasma* infection but not the cyst-forming stage, which resists protocol-specific denaturation. Here, we show that an additional step of enzymatic digestion using mucinase StcE allows full expansion of the *Toxoplasma* cysts, offering a new avenue for a comprehensive examination of the chronic stage of infection using an accessible super-resolution technique.

## Introduction

Expansion Microscopy (ExM) is an imaging protocol that bypasses the diffraction limit of conventional light microscopes (∼200 nm) by physically expanding the biological sample embedded in a gel (1). As an accessible alternative to Electron Microscopy, ExM has been transformative across diverse research fields, allowing nanoscale resolution of protein localisation using light microscopes.

Among the numerous ExM protocols developed so far, Ultrastructure Expansion Microscopy (U-ExM) (2) gained popularity in the field of parasitology due to the improved preservation of subcellular structures (3) U-ExM allows 4-4.5x expansion of the biological specimen (∼90-fold increase in volume) with the preservation of the proteins, in contrast to other approaches which leave only the protein footprint after proteolytic digestion (2, 4). U-ExM therefore enables all-protein staining using non-specific protein dyes like N-hydroxysuccinimde (NHS) esters (5). U-ExM has been widely adopted in Apicomplexan research, generating new insights in to diverse parasite lifecycle stages and clarifying the localisation of over 80 proteins in *Toxoplasma gondii* (6–14), Cryptosporidium parvum *(15, 16)* and *Plasmodium species* (5, 17–26). However, investigations of the chronic *Toxoplasma gondii* cyst have lagged behind.

*Toxoplasma* is a single celled intracellular parasite capable of establishing long-term chronic infection in nearly all warm-blooded animals, including humans (27). *Toxoplasma* has a complex life cycle and differentiates between two non-sexual forms within intermediate hosts (28). During the acute stage of infection, the parasites replicate rapidly as so-called tachyzoites and disseminate around the host’s body. To establish long-term infection, parasites differentiate to slow growing bradyzoites and develop cysts predominantly in the central nervous system and skeletal muscle. Reactivation of latent cysts can cause fatal encephalitis in the immunocompromised or recurrent ocular disease (retinochoroiditis)(29).

*Toxoplasma* tissue cysts are intracellular and are spherical structures 5-100 um in diameter (28), containing a collection of tightly packed bradyzoites. The cyst wall is between 250 and 850 nm thick and composed of a granular layer of proteins and carbohydrates underneath the limiting membrane (a modified version of the parasitophorous vacuole (PV) membrane which forms as the parasite invades the host cell). The accumulation of glycoproteins in the cyst wall ensures structural and chemical rigidity (30), and is thought to allow parasite transmission to infect a new host following ingestion of undercooked meat.

*Toxoplasma* cysts can also be generated *in vitro* in tissue culture, which has proved a valuable model for studying cyst biology. *Toxoplasma* tachyzoites differentiate to bradyzoites under stress conditions with alkaline stress the most commonly used method (31, 32). However, recent advances have improved the model with spontaneous differentiation to bradyzoites in myotubes and neurons, producing longer lasting cysts (14-28 days) (33, 34).

The cyst wall contains markers exclusively expressed during the chronic stage (35). One of the early chronic-stage markers, CST1, is major structural element of the cyst wall and parasites lacking *cst1* form fragile cysts (30). CST1 has a highly O-glycosylated mucin domain which binds to the lectin Dolichos Biflorus agglutinin (DBA) (30, 36). The structure of CST1’s mucin domain includes 20 threonine-rich tandem repeats, which undergo O-GalNAc glycosylation. This glycosylation process is initiated by a group of enzymes known as polypeptide N-acetylgalactosaminyltransferases (ppGalNAc-Ts, (30, 36)). Previous studies using electron microscopy have shown that CST1 is expressed in the granular material in the cyst wall under the limiting membrane (37). It’s been hypothesised that the high glycosylation of the CST1 mucin domain could act as a bonding agent for other proteins associated with the cyst wall, such as dense granule proteins (GRAs) (38).

*Toxoplasma* GRAs are secreted from dense granules into the PV lumen or out into the host cell and perform a variety of functions during infection including mediating host–parasite interactions, modification of the PV, and establishment of intravacuolar network (IVN) of highly curved nanotubules (39). The latter is involved in connecting tachyzoites in the PV and in the salvage of lipids (40, 41) and cytosolic proteins (42) from the host. Some of the GRA proteins, especially those associated with the IVN network (GRA2, GRA6, GRA4, and GRA12), relocalize to the forming cyst wall and impact CST1 localisation (38). Of these, GRA2 is essential for IVN formation (43), acute virulence and cyst formation (44, 45), yet details of its function during the chronic stage remain unclear. GRA2 appears to have a dynamic cyst wall localization being present in early cysts at day 7 (D7) and D10, with its location overlapping with the CST1 signal (38), while absent from the wall on late cysts (46).

Here, we apply U-ExM to *Toxoplasma* tissue cysts produced *in vitro* demonstrating that the original protocols do not successfully expand these structures. This problem is solved by the addition of an enzymatic digestion step with mucin-selective protease StcE (47), during the sample preparation. StcE cuts through the mucin domains of the major structural protein of the cyst wall CST1, allowing the cyst to expand fully, without damaging CST1 or other protein epitopes. We used this method to assess the co-localisation of CST1 and GRA2 in the cyst wall and observed distinct localisation at this higher resolution. The addition of the StcE step to the U-ExM protocol opens new avenues to precisely localize proteins in bradyzoites, as exemplified here with the proteins associated with the cyst wall, to aid in understanding their role in cyst formation and growth.

## Materials and methods

### Cell culture

Primary human foreskin fibroblasts (HFFs) (ATCC) were maintained in Dulbecco’s modified Eagle’s medium (DMEM, Sigma-Aldrich) with 4.5 g/L glucose, 25 mm HEPES and 1% v/v GlutaMAX (Gibco) supplemented with 10% v/v fetal bovine serum (FBS, Gibco) at 37°C with 5% CO2.

### Parasite strains and culture

*Toxoplasma* gondii ME49Δ*ku80*Δ*hxgprt (48)* and 76K tachyzoites were propagated *in vitro* in HFFs using DMEM supplemented with 2% v/v heat-inactivated FBS, 1% v/v glutaMAX, and 1% v/v penicillin-streptomycin solution (Gibco). Tachyzoites were grown in ventilated tissue culture flasks at 37°C and 5 % CO2. Prior to infection, intracellular parasites were isolated by syringe passage with 23-gauge blunt needles (SAI Infusion Technologies) and filtration through a 5 μm membrane filter (Sartorius).

### Parasite infections

For confocal and U-ExM imaging, HFFs were seeded on 13-mm no. 1.5 coverslips (SLS) in 24-well plates. Confluent monolayers at least 7 days old were infected at a multiplicity of infection (MOI) of 1 for 24 hours. Cells were fixed in 4% formaldehyde (FA, Thermo Scientific) for 15 min at RT.

### Conversion to bradyzoites

Bradyzoite cultures were obtained by infecting HFF monolayers coverslips with tachyzoites at an MOI of 1 for 3.5 hours, followed by media change to filtered RPMI 1640 media (Sigma-Aldrich) supplemented with 1% v/v FBS, and 50mM HEPES (Sigma-Aldrich) and brought to pH 8.2. The bradyzoite culture was grown with ambient CO2 at 37°C for 7 days with daily media changes. Cells were fixed in 4% FA for 15 min at RT.

*T. gondii* infection of primary neuronal culture obtained from the hippocampus of postnatal rats was performed as described previously (33). Briefly, after the dissection of the brains, hippocampi were mechanically dissociated and resuspended in Neurobasal A, a medium supplemented with GlutaMAX and B27 neural supplement with antioxidants (Gibco). Cells were plated at a density of 100 000 cells/cm^2^ in poly-L-lysine coated 24-well plates containing coverslips. Brain cells were grown for 14 days before infection. Each well was infected by 1.10^4^ tachyzoites of the 76K strain resuspended in the Neurobasal A medium. The culture was grown for an extra 14 days to obtain fully mature cysts as described in Mouveaux *et al*.

### Standard immunofluorescence assays

For immunofluorescence assays, fixed cells were permeabilised with 0.1% Triton X-100 (Sigma-Aldrich) in PBS for 2 min at RT for tachyzoites or 0.2% Triton X-100, 0.1% glycine (Sigma-Aldrich), and 0.2% BSA in PBS for 20 min on ice for bradyzoites. Samples were then incubated in blocking solution (2% BSA in PBS) for 1 hour, followed by 45 min incubation with antibodies/stains (diluted in blocking solution).

Infected cells, depending on experiment, were stained using antibodies and dyes listed in **Table 1**. Coverslips were washed three times after primary and secondary antibody/stain incubations, and coverslips were subsequently washed and mounted with Vectashield Antifade softset mountant (H-1000; Vector Laboratories). Parasites were visualized on a Zeiss LSM980 laser confocal microscope.

**Table 1.** Key resources table.

| Category | Reagents | Manufacturer | Additional information |
| --- | --- | --- | --- |
| U-ExM protocol reagents | Acrylamide, 40% (AA) | Sigma-Aldrich |  |
|  | Ammonium persulfate (APS) | Sigma-Aldrich |  |
|  | Formaldehyde, 37% (FA) | Sigma-Aldrich |  |
|  | N,N'-methylenebisacrylamide, 2% (Bis) | Sigma-Aldrich |  |
|  | Sodium Chloride (NaCl) | Thermo Fisher |  |
|  | SDS, 20% | ThermoFisher |  |
|  | Sodium Acrylate (SA) | AKSci |  |
|  | TEMED | Sigma-Aldrich |  |
|  | Tris base | ThermoFisher |  |
|  | Tween20 | Sigma-Aldrich |  |
|  | Poly-D-lysine | Thermo Scientific |  |
|  | StcE mucinase, 20 uM | Gift from Kayvon Pedram, (50) | Commercially available from Merk, Cat N SAE0202 |
| Dyes, stains, antibodies, and mountants | DAPI (4',6 Diamidino 2 Phenylindole, Dilactate) | Invitrogen | 1:1000 (std), 1:100 (ExM) |
|  | Dolichos Biflorus Agglutinin (DBA), biotinylated | Vector Laboratories | 1:2500 (std), 1:1000 (ExM) |
|  | anti-GRA2 mouse | Biotem | 1:1000 (std), 1:300 (ExM) |
|  | anti-SAG1 rabbit | Abcam | 1:1000 (std) |
|  | anti-CST1 rabbit | Gift from L. Weiss, (30) | 1:200 (std), 1:100 (ExM) |
|  | anti-TgGAP45 rabbit | Gift from D. Soldati-Favre, (52) | 1:10000 (std), 1:7000 (ExM) |
|  | AlexaFluor 488 goat anti-mouse superclonal | Thermo Scientific | 1:1000 (std), 1:300 (ExM) |
|  | AlexaFluor 568 goat anti-mouse | Thermo Scientific | 1:1000 (std), 1:300 (ExM) |
|  | AlexaFluor 488 goat anti-rabbit | Thermo Scientific | 1:1000 (std), 1:300 (ExM) |
|  | AlexaFluor 568 goat anti-rabbit | Thermo Scientific | 1:1000 (std), 1:300 (ExM) |
|  | Atto 647N goat anti-rabbit | Sigma-Aldrich | 1:1000 (std), 1:300 (ExM) |
|  | NHS ester, Atto 565 conjugate | BioReagent | 1:300 (std), 1:300 (ExM) |
|  | Streptavidin, AlexaFluor 488 conjugate | Invitrogen | 1:1000 (std), 1:300 (ExM) |
|  | Streptavidin, AlexaFluor 568 conjugate | Invitrogen | 1:1000 (std), 1:300 (ExM) |
|  | Mounting media VECTASHIELD Antifade | Vector Laboratories |  |
| Tissue culture media and reagents | Dulbecco's modified Eagle's medium | Thermo Fisher |  |
|  | RPMI 1640 media | Sigma-Aldrich |  |
|  | Fetal bovine serum | Thermo Fisher |  |
|  | Bovine serum albumin | Sigma-Aldrich |  |
|  | HEPES | Sigma-Aldrich |  |
|  | Penicillin-streptomycin solution | Thermo Fisher |  |
|  | GlutaMAX | Thermo Fisher |  |
|  | Formaldehyde, 16% | ThermoFisher |  |
|  | Glycine | Sigma-Aldrich |  |
|  | Triton X-100 | Sigma-Aldrich |  |
| Tissue culture and staining consumables | Blunt needles 23G | SAI Infusion Technologies |  |
|  | Cellulose Acetate membrane filter, 5 µm | Sartorius |  |
|  | Coverslips No 1.5, Dia 13 mm | SLS |  |
|  | Press-to-seal silicone isolator (13mm*0.8mm) | Sigma-Aldrich |  |
|  | 35-mm high glass-bottom petri µ-dishes | Ibidi |  |

### Ultrastructure expansion microscopy (U-ExM)

Gambarotto and colleagues published a detailed U-ExM protocol in 2021 (49). For reagents list, see **Table 1**. Briefly, fixed infected HFFs or primary neuronal cultures on coverslips were first permeabilised (tachyzoites: 0.1% Triton X-100 in PBS for 2 min at RT; bradyzoites: 0.2% Triton X-100, 0.1% glycine, and 0.2% bovine serum albumin (BSA) in PBS for 20 min on ice). They were then incubated in protein crosslinking prevention solution (final conc. 1.4% PFA/2% Acrylamide (AA)) at 37°C with no shaking overnight.

To obtain fully expanded cysts, protein crosslinking prevention was followed by the incubation with 100nM StcE mucinase in PBS (gift from Kayvon Pedram, (50)) for 4 hours at 37°C with no shaking. StcE is also commercially available (SAE0202, Merk). Coverslips were briefly washed 3x PBS and prepared for gelation.

The gelation solution consisted of a monomer solution, topped up with TEMED (Sigma-Aldrich) and APS (Sigma-Aldrich) solutions just before the sample application (**Table 2**). All components except PBS were stored at -20°C and kept on ice just before the application.

**Table 2.**
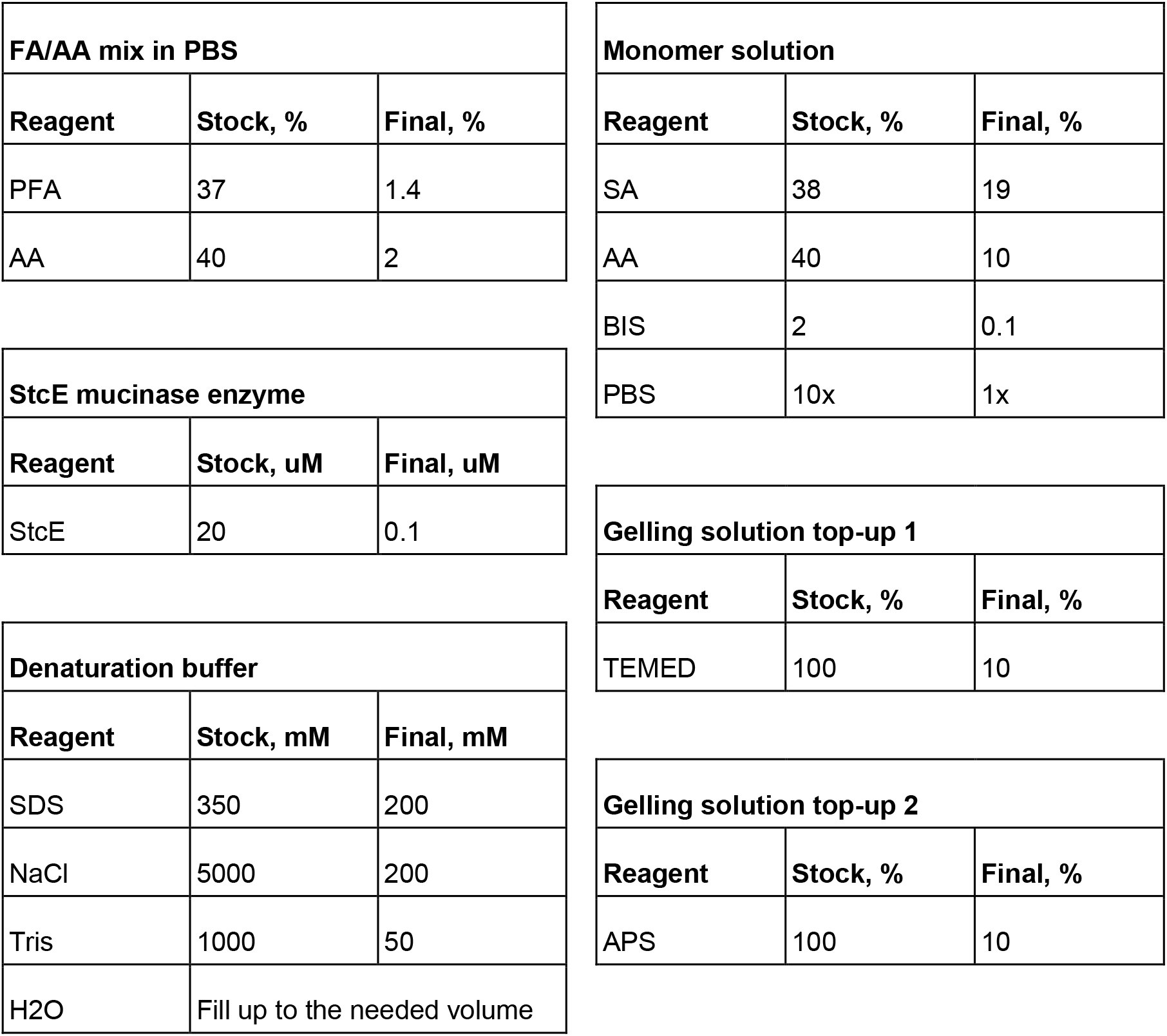
Expansion microscopy buffers.

The gelation chamber was adapted from (51) and was prepared as follows. A clean press-to-seal silicone isolator (13mm diameter x 0.8mm depth, JTR13R-1.0, Sigma-Aldrich) was mounted on a parafilm-wrapped glass slide. The device was put on a wetted piece of white roll paper, stretched in a plastic chamber (eg. a 24-well plate lid), placed for 5 minutes in -20°C freezer and then kept on ice.

120 μL of fresh gelation solution was placed into the well created by silicon isolator. Immediately after, the sample coverslip was dabbed from excess PBS and placed cells down on the top of the well. The gelation chamber was left for 5 min on ice to aid monomer permeation, and then placed into 37°C incubator for 1 hour for polymerisation.

Once polymerised, the silicone isolator was peeled off, coverslips lifted from the parafilm-covered glass using tweezers, put in 6-well plate (1 gel/well), topped with 2mL denaturation solution (**Table 2**) and left for 15 min on a rocker. By the end of 15 min, gels that were detached from the coverslips were placed in in a 1.5mL Eppendorf tube with the fresh denaturation solution and incubated for 1.5 hours at 95°C in a heating block with no shaking. If gels were not detached by that point, they were left for another 5-10 min on the rocker. After denaturation, gels were then expanded in 250-300mL ddH2O overnight.

The next day, gels underwent 2x15min incubation in PBS (no shaking) to shrink in size, and then cut to the needed shape using scalpel blade (to fit a well in 24-well plate). Note that the gels shrink further during the staining process. Gels were then blocked for 1 hour in a blocking solution (2% BSA in PBS) at 37°C while shaking, and then incubated with the primary and secondary antibodies or stains for 2.5 hours in the same conditions, with 3x10min washes in PBS-0.1%Tween20 solution on a rocker at RT in between incubation steps.

Infected cells, depending on experiment, were stained using antibodies and dyes listed in **Table 1**. Parasites were visualized on a Nikon Ti2 CSU-W1 Spinning Disk microscope or a Zeiss LSM980 laser confocal microscope in 35-mm high glass-bottom petri μ-dishes (Ibidi), covered in-house with Poly-D-lysine (Thermo Scientific) for 1h, followed by 3x ddH2O washes, and then dried.

### Image acquisition

Imaging was performed in Centre Optical Instrumentation Laboratory (COIL), University of Edinburgh. Two microscopes were used to visualise expanded samples: Nikon Ti2 CSU-W1 Spinning Disk Confocal (objective: 100x: Plan Apo TIRF, Oil, 1.45 NA) and Zeiss LSM980 laser-scanning confocal microscope (objectives: 20x: Plan Apochromat, Air, 0.8 NA; 100x: Alpha Plan Apochromat, Oil, 1.45 NA, DIC). Initially Nikon Ti2 CSU-W1 Spinning Disk Confocal was used for faster gel imaging to avoid potential gel drift. However, poly-D-lysine covering substantially reduced the drift when imaged on more sensitive Zeiss LSM980 laser-scanning confocal microscope, and thus, all consequent imaging was performed using Zeiss.

### Image Processing and Data Analysis

The images were exported using either Zen Blue software (Zeiss) or ImageJ. The brightness was adjusted for display purposes, but the measurements were taken on raw unprocessed files.

The expansion factor was determined by the comparison of the average cross-section length of parasite nuclei stained with DAPI nuclear stain (along the longest axis) in the tachyzoites before and after expansion. Data analysis and statistical tests (t-tests) were performed in GraphPad Prism software.

## Results

### Standard Ultrastructure Expansion Microscopy protocol does not fully expand *T. gondii* cysts

Standard U-ExM protocols have been successfully applied to extracellular *Toxoplasma* and tachyzoites within infected cells (reviewed in (3)). To assess whether this method could be applied to *Toxoplasma* cysts, we first compared U-ExM protocol on *Toxoplasma* infected cells grown under standard conditions (tachyzoites) or under pH stress to induce conversion to bradyzoites.

The standard U-ExM protocol of expansion *T. gondii* parasites from a monolayer of infected cells takes three days (**Fig.1A**) and consists of the following steps: fixation and permeabilisation of the tissue, incubation in crosslinking preventative and protein anchoring solution, gelation and polymerisation, followed by denaturation at 95°C with 200mM SDS, antibody labelling and final expansion of the gelled sample in water (2, 49).

For the tachyzoite samples, the expansion was successful (**Fig.1B-E**) with an average expansion factor of 4.3x (**Fig.1C**), similar to the published values of 4-4.3x for *T. gondii* (10, 23, 53) *Plasmodium falciparum*, and *Cryptosporidium parvum parasites* (3).

NHS ester binds all proteins, and its fluorophore-conjugated version was used to visualise the cell architecture, which demonstrated that the host-cell environment tightly surrounds the PV in the expanded sample, leaving no gap (**Fig.1E, white arrows**). In contrast, when comparing the expanded and non-expanded *T. gondii* cysts of D7 bradyzoite culture with anti-CST1 antibody used as a cyst wall marker (**Fig.1F-G**), a large black gap was consistently visible between the cyst and the surrounding tissue of the expanded host cell (**Fig. 1G**). Furthermore, quantification of the bradyzoite nuclei cross-section before and after expansion revealed a smaller 2.5x expansion factor compared to 4.3x in tachyzoite samples (**Fig.1C**), rendering the standard U-ExM protocol insufficient for super-resolution studies of the *Toxoplasma* cyst wall.

**Figure 1.**
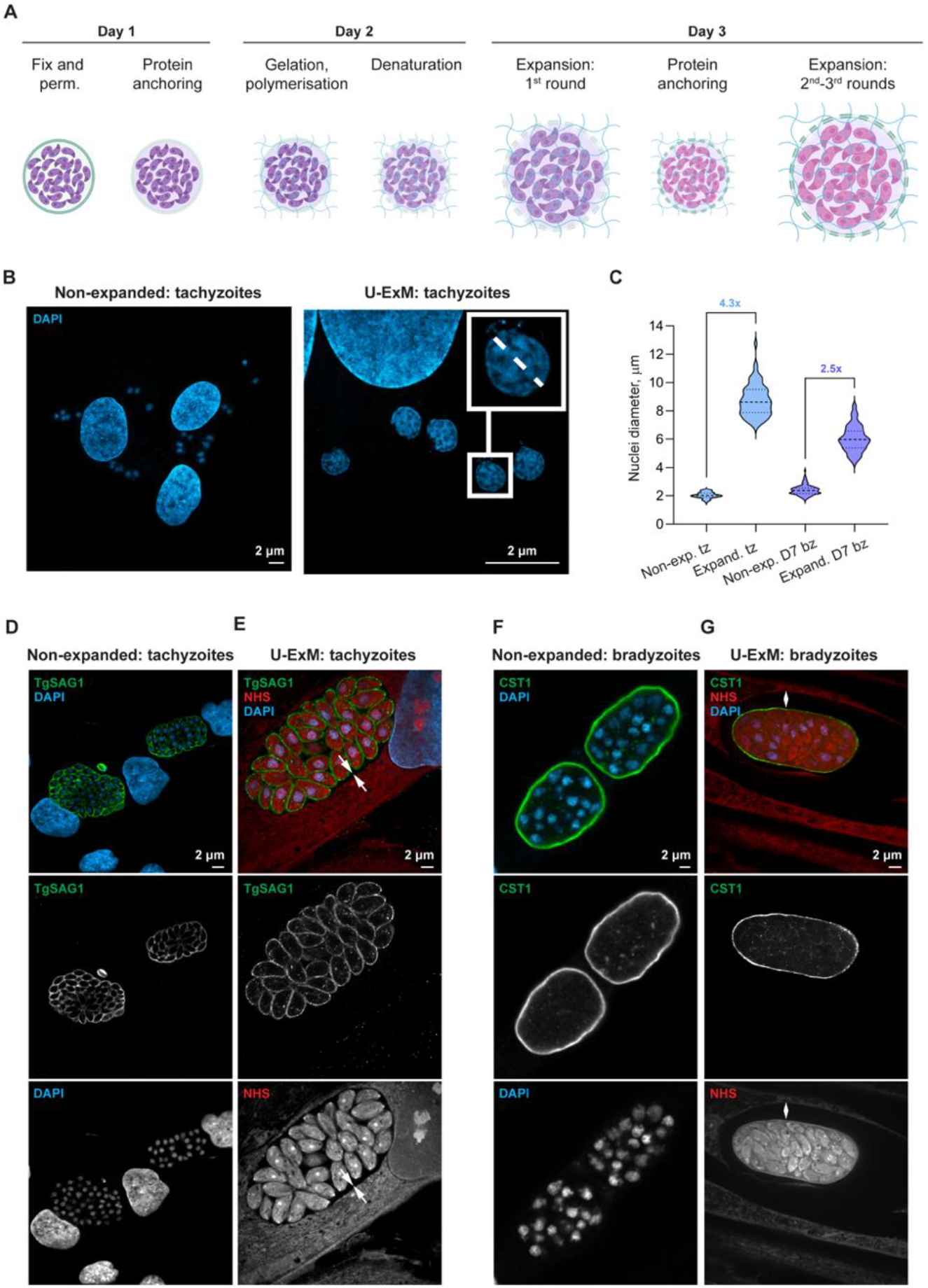
Standard U-ExM protocol expands intracellular *T. gondii* tachyzoites but fails to expand the cyst wall. **(A)** An overview of the standard U-ExM protocol workflow (created with BioRender.com). **(B)** Confocal images of HFF monolayer infected with Type II ME49Δku80Δhxgprt tachyzoites, fixed 24h pi, and labelled with DAPI nuclear stain. Large nuclei belong to host cells, smaller are tachyzoite nuclei. Insert shows the example of tachyzoite nuclei measurement along its longest diameter to calculate the expansion factor of the U-ExM sample. **(C)** The expansion factor was determined by comparison of the average cross-section of tachyzoite (light blue) and bradyzoite (violet) nuclei stained with DAPI in non-expanded samples versus U-ExM-processed. The tachyzoite expansion factor is 4.3x, while only 2.5x for bradyzoites (for both unpaired t-test, p <0.0001) (n_non-exp. tz_= 79, n_exp. tz_=131, n_non-exp. bz_=201, n_exp. bz_=112). **(D**,**E)** Immunofluorescence images of HFFs infected with ME49Δku80Δhxgprt tachyzoites for 24h. Confocal images (single optical sections) of non-expanded tachyzoites in (D) versus U-ExM-processed shown in (E), both probed with anti-TgSAG1 antibody (green) and DAPI (blue), while U-ExM-processed samples were also stained with all-protein NHS-565 dye (red). White arrows indicate continuous host-cell environment surrounding the parasitophorous vacuole. **(F**,**G)** Immunofluorescence images of ME49Δku80Δhxgprt infected HFFs grown under bradyzoite inducing conditions for 7 days. Confocal images (single optical sections) of non-expanded D7 cysts in (F), versus U-ExM-processed shown in (G). Both samples were probed with anti-CST1 antibody (green) and DAPI (blue), while U-ExM-processed samples were also stained with all-protein conjugated NHS-565 dye (red). White arrows show a gap between the fully expanded host cell and only partially expanded cyst. Tz – tachyzoites, Bz – bradyzoites, exp. – expanded.

### Incubation with the mucin-selective protease StcE prior U-ExM processing allows full expansion of the *T. gondii* cyst in fibroblasts and primary neurons

It is well established that the *Toxoplasma* cyst wall consists of an enrichment of glycoproteins including CST1, SRS13, and TgPPG (30, 54, 55). Of these, CST1 is a major structural component of the cyst, and is a SAG1-related sequence protein with a heavily glycosylated mucin domain (30). We initially addressed the problem of partially expanding cysts by attempting enzymatic digestion using Proteinase K (ProK), commonly used in protein retention expansion microscopy (pro-ExM), alongside the denaturation step (data not shown). However, carbohydrate modifications of the cyst wall were resistant to this type of proteolytic digestion. As an alternative enzymatic approach that would disrupt densely packed glycosylated domains without reducing protein signals, we included a mucinase StcE step in the protocol. StcE is a mucin-selective protease with a specific peptide- and glycan-based cleavage motif that digests the mucin domain, as shown in **Figure 2A,B** (47). We first tested whether the StcE enzyme affects the proteins associated with the cyst wall in a non-expanded sample (**Fig. 2C**) by introducing a StcE enzymatic treatment after the permeabilisation step. Cysts from the D7 bradyzoite culture labelled with antibodies against CST1 and GRA2 show the expected protein localisations after 4h incubation with StcE at 37°C, showing that it doesn’t destroy the cyst wall. In fact, both signals increased in intensity following the StcE step, possibly as a result of increased antibody accessibility.

**Figure 2.**
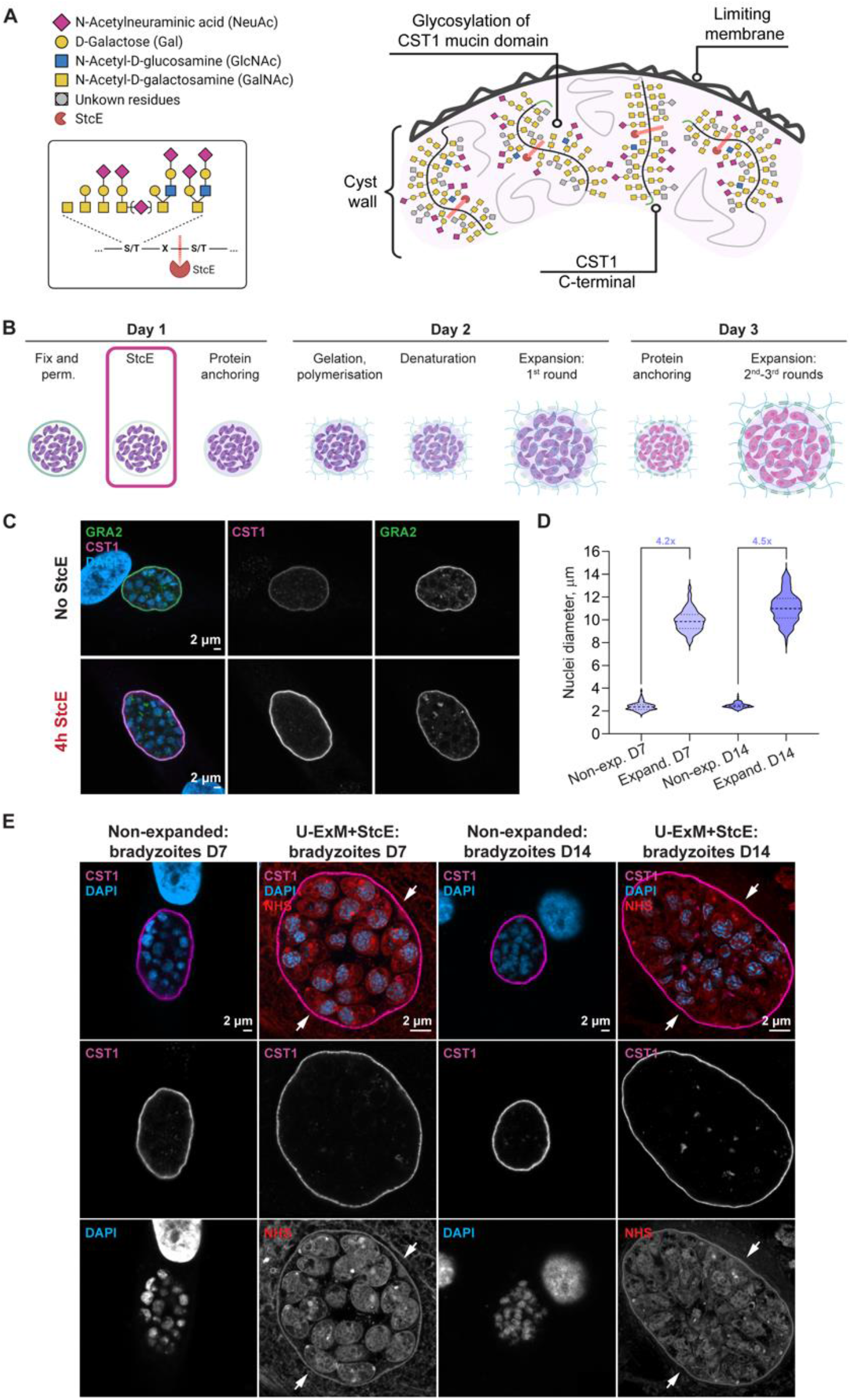
U-ExM protocol complemented with StcE digestion fully expands the cyst wall of D7 and D14 cysts. **(A)** Proposed mechanism of mucin domain cleavage by StcE mucin-selective protease. Diagram shows part of the cyst wall, depicting the limiting membrane (faces host-cell environment), and the CST1-positive layer with the ribbon-like glycoprotein structures of CST1 mucin domains. The glycoprotein structures illustrated in the cyst wall diagram correspond to those known to be associated with CST1 (featuring GalNAc-GalNAc as a DBA-binding site (30, 36)), and those that are preferred cleavage targets of StcE. Insert depicts the StcE cleavage point on the glycopeptide chain. StcE cleaves before the second S/T within the motif S/T*–X–S/T, where the asterisk indicates modification with a glycan and X can be any amino acid or absent (47). **(B)** Adapted U-ExM protocol workflow with added StcE enzymatic treatment (100 nM, 4h, 37°C, no shaking) after the permeabilization step. **(C)** Representative confocal images (single optical sections) of non-expanded HFF monolayer infected with Type II ME49Δku80Δhxgprt D7 cysts with and without StcE treatment. Samples were labelled with antibodies against GRA2 (green), CST1 (magenta), and stained with DAPI nuclear stain, followed by imaging under the identical acquisition settings. *Top row*: no StcE treatment (control); *bottom row*: samples treated with 4h StcE at 37°C after permeabilization step. **(D)** The expansion factor of 4.2x was determined by the comparison of the average cross-section length of D7 bradyzoite nuclei in CST1-labelled cysts stained with DAPI in non-expanded HFF samples versus U-ExM+StcE-processed HFFs (unpaired t-test, p <0.0001), and 4.5x in non-expanded neurons versus U-ExtM+StcE processed neurons (unpaired t-test, p <0.0001) (n_non-exp.bz HFFs_= 201, n_non-exp.bz neurons_= 218, n_exp.bz HFFs_=185, n_exp.bz neurons_=123). **(E)** Confocal images (single optical sections) of non-expanded D7 (in HFFs) and D14 cysts (in neurons) versus expanded using U-ExM+StcE. All samples were probed with anti-CST1 (magenta) antibody and DAPI (blue), with the additional all-protein NHS-565 stain (red) for U-ExM+StcE. White arrows indicate continuous host-cell environment surrounding cyst wall. Tz – tachyzoites, Bz – bradyzoites, exp. – expanded. Diagrams in panels (A) and (B) created with BioRender.com.

We then introduced StcE treatment after the permeabilization step in the U-ExM protocol. With this optimised protocol we observed successful expansion of 7-day *Toxoplasma* cysts with an expansion factor of 4.x **(Fig. 2D)**, similar to that of tachyzoites. Crucially, no gap is observed between the expanded cyst and the host cell environment (**white arrows, Fig. 2E)**.

It was recently shown by Mouveaux et al., that D14 *in vitro* cysts were able to infect a mouse following oral gavage, while infection with D7 cysts was unsuccessful (33). This may be that the cyst wall continues to mature and becomes more robust over time, or that the parasites differentiate further. In order to verify the StcE enhanced U-ExM protocol on infectious cysts, we tested 14-day old cysts generated in primary neuronal cultures from postnatal rat hippocampus alongside with the D7 cysts from the HFF monolayer. Reassuringly, a similar expansion factor was observed **(Fig. 2D)**, with no gap between the packed cyst structure and the surrounding cell **(Fig. 2E)**. To further assess the method, expanded *in vitro* cysts were probed with antibodies against the inner membrane complex, GAP45, and the secreted protein GRA2 along with anti-CST1 for the cyst wall **(Fig. 3A-D)**. All antibodies showed their expected localisation patterns, with GAP45 outlining the parasite shape, and GRA2 split between intracellular staining within dense granules and localised around the periphery of the cyst.

**Figure 3.**
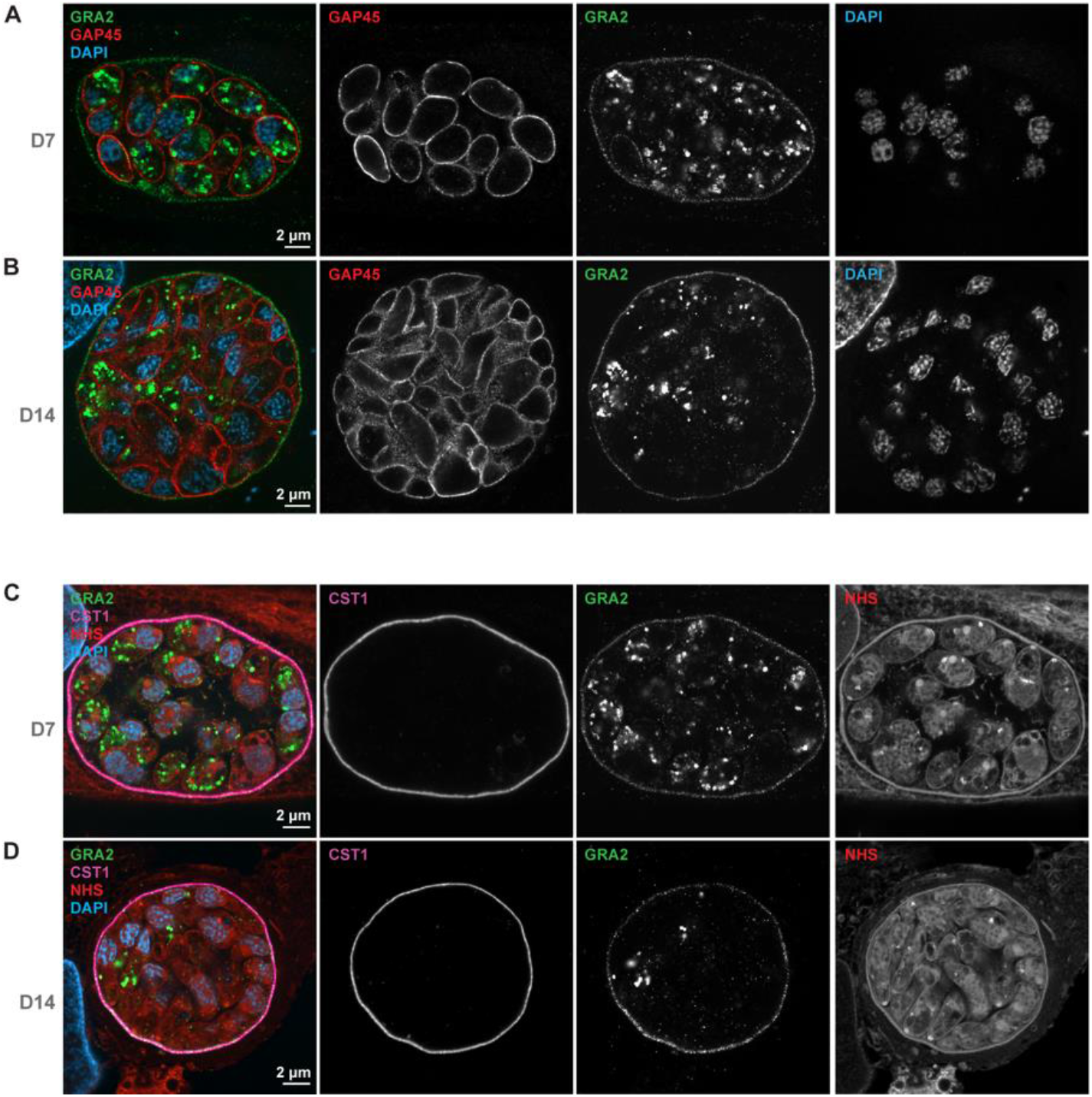
U-ExM+StcE protocol preserves protein signals in *T. gondii* cysts *in vitro* (D7 fibroblasts, D14 neurons) **(A-B)** Confocal image (single optical section) of the U-ExM+StcE-expanded HFF monolayer with Type II ME49Δku80Δhxgprt D7 cyst **(A)** and primary neuronal culture (derived from P0 rat hippocampus) with Type II 76K D14 **(B)** cyst at 100x magnification. Samples were labelled with antibodies against GRA2 (green), GAP45 (magenta), biotinylated DBA (not shown), and stained with DAPI nuclear stain. Each row consists of a merged image on the left, followed by separate channels in grey. **(C-D)** Confocal image (single optical section) of the U-ExM+StcE-expanded HFF monolayer with Type II ME49Δku80Δhxgprt D7 cyst **(C)** and primary neuronal culture (derived from P0 rat hippocampus) with Type II 76K D14 **(D)** cyst at 100x magnification. Samples were labelled with antibodies against GRA2 (green), CST1 (magenta), and stained with NHS-565 and DAPI.

### Modified U-ExM-StcE protocol reveals that GRA2 is only partially localised to the CST1-positive layer in the cyst wall

While many proteins secreted from the dense granules are known to localise to the cyst wall (46, 56, 57), current microscopy techniques have not allowed more detailed visualisation of substructures. We therefore used the U-ExM+StcE protocol to clarify the location of GRA2 relative to CST1 protein in the *T. gondii* cyst wall in HFFs (D7) and primary rat neuronal cultures (D14). While previous data has shown smooth overlapping signal for CST1 and GRA2 in *in vitro* cysts (38), we observe a more granular distribution of GRA2 in the cyst wall compared to the smooth signal of CST1 with StcE enhanced U-ExM (**Fig.4A**). The GRA2 granules in the expanded cyst are observed on both the outer and inner sides of the CST1 layer, with enrichments visible on the inner side of the wall (**Fig. 4B-E**). In viewing the z-stack of images through the wall these GRA2 structures appear to span through the CST1-positive layer **(Supp Movie 1)**. A similar pattern was observed in D14 cysts from infected primary neuronal cultures (**Fig.4 F-J**).

**Figure 4.**
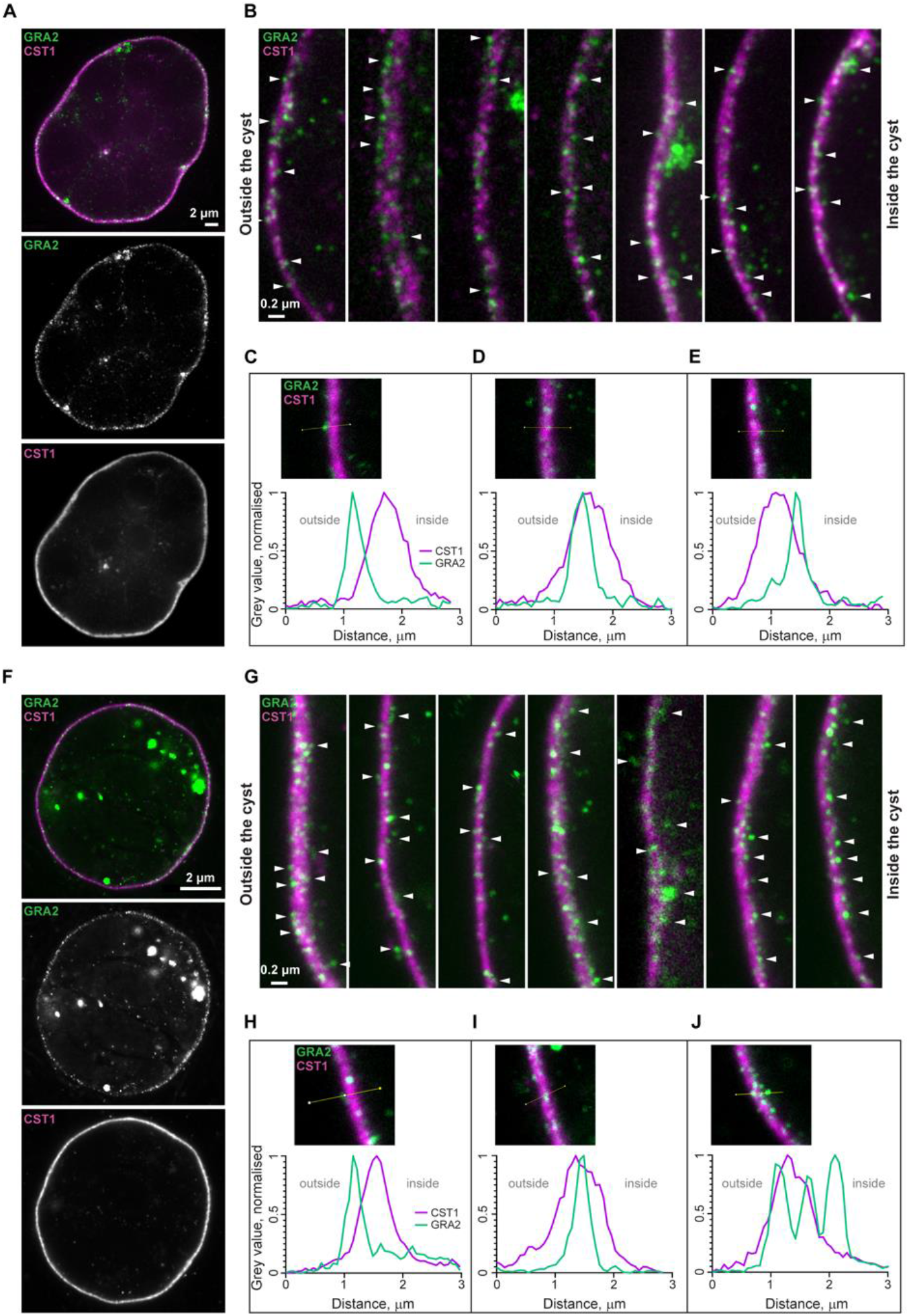
U-ExM of *in vitro* cysts shows GRA2 spanning the CST1-positive layer in HFF D7 and neuronal D14 *T. gondii* culture. **(A)** Confocal image of the U-ExM+StcE-expanded HFF monolayer with Type II ME49Δku80Δhxgprt D7 cyst at 100x magnification (single optical section of the widest cross-section of the cyst). Sample wase labelled with antibodies against GRA2 (green) and CST1 (magenta). **(B)** Zoomed cyst-wall regions of the three D7 cysts. White arrows highlight examples GRA2 puncta inside and outside the CST1 layer. **(C-E)** Fluorescence intensity profiles showing different locations of GRA2 relative to CST1 in a single U-ExM+StcE-expanded D7 cyst grown in HFF monolayer. Normalised intensity levels (grey values) obtained from the raw images were plotted against the length of the sampled area (yellow line, image insert). **(C)** GRA2 localises outside the CST1 layer, on the exterior side of the cyst wall, facing host-cell environment. **(D)** GRA2 fully co-localises with CST1. **(E)** GRA2 localises outside the CST1 layer, on the interior side of the cyst wall facing cyst matrix. **(F)** Confocal image of the U-ExM+StcE-expanded primary neuronal culture (derived from P0 rat hippocampus) with Type II 76K D14 cyst at 100x magnification (single optical section of the widest cross-section of the cyst), presented as in (A). **(G)** Zoomed cyst-wall regions of the three D14 cysts. White arrows show GRA2 puncta inside and outside the CST1 layer. See also Supp Movie 1. **(H-J)** Fluorescence intensity profiles showing different locations of GRA2 relative to CST1 in a single U-ExM+StcE-expanded Type II 76K D14 cyst grown in a primary neuronal culture. Normalised intensity levels (grey values) obtained from the raw images were plotted against the length of the sampled area (yellow line, image insert). **(H)** GRA2 localises outside the CST1 layer, on the exterior side of the cyst wall facing host cell environment. **(I)** GRA2 fully co-localises with the CST1. **(J)** Three GRA2 punctae localise across the CST1 layer: one on the exterior side of the cyst wall, one in the middle of CST1-positive layer, and one outside, facing cyst matrix.

## Discussion

Despite the importance of chronic *Toxoplasma* cysts in transmission and their reactivation in human disease, the detailed structure of the cyst is yet to be clarified. While EM has provided key insights into the wall morphology, resolution has limited the ability to precisely localise many cyst wall proteins. A relatively new modification of the sample expansion technique, U-ExM, is an easy, cost-effective alternative to the established super-resolution approaches, allowing direct mapping of the sample morphology to protein location.

When attempting to use U-ExM for cyst wall protein localisation, we discovered that the standard U-ExM protocol fails to expand the cyst wall up to the expected 4-4.3x expansion factor. The addition of the enzymatic step targeting glycosylated mucin domains of CST1 and, potentially, SRS13 proteins using StcE mucinase solves the ineffective expansion, allowing protein localisation studies in fully expanded cysts. We confirmed our results in Day 7 and Day 14 cysts grown in HFF monolayers and primary rat neuronal cell culture, respectively.

We then used the technique to clarify the location of the GRA2 protein relative to the CST1-positive layer in the cyst wall. Previously published studies based on standard confocal images show full co-localisation of GRA2 and CST1 both appearing as a smooth signal across the cyst wall (38). Using modified U-ExM+StcE protocol, we show that GRA2 is distributed as puncta across the still smooth CST1-positive layer, which was consistent in both in D7 *Toxoplasma* cysts in fibroblasts and and D14 cysts in neurons *in vitro*. Moreover, fully expanded cysts show GRA2 spanning the CST1-positive layer, providing previously unknown details about the GRA2 location in the *Toxoplasma* cyst. Enrichments of GRA2 signal were observed underneath the CST1 layer, with putative spirals going through the CST1 layer. In the future, it will be interesting to investigate if these represent clusters of tubular network of the IVN and colocalise with other GRAs.

Early work on GRA2 localisation using immuno-electron microscopy with labelled golden particles, showed GRA2 secretion from the tachyzoites into the PV lumen and its association with the IVN, although the GRA2 localization doesn’t follow any specific pattern (58). It is well known that GRA2 is essential for the correct organisation of the IVN in tachyzoite PVs (43), and impacts the cyst wall and matrix during the chronic stage (38). This, and its relocation to the cyst wall with the subsequent co-localisation with the major structural protein CST1, suggests that GRA2 retains its role in the organisation of the parasitic structures after the stage conversion, however, its precise function is still unclear.

Although a significant portion of GRA2 relocalises to the cyst wall, some remain in the cyst matrix as puncta (38). Regular (non-expansion) immunofluorescence assays depict cyst-wall-associated GRA2 as a smooth continuous signal. We show that GRA2 retains its puncta appearance in the cyst matrix and the cyst wall.

Despite the growing number of proteins localised to the cyst wall over the past few years (56, 57, 59), we still don’t know their functional relevance in the development and maturation of the cyst. By revealing more precise protein localisations, U-ExM+StcE can play a key role in investigating their role by expanding our knowledge of substructures within the wall. Indeed, with the expansion of *Toxoplasma* cysts over time, and their subsequent rupture to release parasites, it is clear that the cyst wall is subject to remodelling.

In conclusion, the U-ExM+StcE protocol is a powerful addition to the microscopy toolbox aimed at the investigation of *in vitro Toxoplasma* cysts with nanoscale resolution, joining other recent efforts to adapt expansion protocols for non-mammalian samples via tailored enzymatic disruption (60). In our case, the simplicity of a single-enzyme treatment with a pan-mucinase (50) to break down the mucin-rich glycocalyx of the *Toxoplasma* cyst wall underscores the importance of biochemical characterization of structural biomolecules for successful Expansion Microscopy.

## Acknowledgments

We thank all members of the Young lab as well as Nisha Philip and Joanne Thompson for critical discussion and support. We particularly thank Johanna Dornell for technical assistance on HFF culture. We thank Paul Tillberg (Janelia Research Campus) for advice on enzymatic digestion and for referring us to Kayvon Pedram. We also thank Dominique Soldati-Favre (University of Geneva, Switzerland) for providing anti-GAP45 antibody, and Louis Weiss (Albert Einstein College of Medicine, USA) for providing anti-CST1 antibody. This work was funded by an MRC Career Development award (MR/V03314X/1) supporting JCY and KB. MG is a CNRS researcher. MG and FL were supported by a Transdisciplinary Research Projects grant from the Pasteur Institute in Lille awarded to MG. Imaging was performed in Centre Optical Instrumentation Laboratory (COIL), which is supported by funding for the Wellcome Discovery Research Platform for Hidden Cell Biology [226791] and we gratefully acknowledge support from the Microscopy core, particularly David Kelly and Toni Mchugh for training and technical assistance. The use of Nikon Ti2 CSU-W1 Spinning Disk and Zeiss LSM980 confocal microscopes was supported by the Wellcome Trust grant [203149].

## Data availability

All data is freely available upon request.

## Open Access Statement

For the purpose of open access, the author has applied a ‘Creative Commons Attribution (CC BY) licence to any Author Accepted Manuscript version arising from this submission.

## Supplementary data

### Supplementary movie 1

#### The GRA2 granules in the expanded cyst span through the CST1-positive layer

ME49Δku80ΔhxgprtME49 bradyzoite cysts grown in HFFs for 7 days, probed with antibodies against GRA2 (green) and CST1 (magenta), and expanded using UExM+StcE protocol as in Figure 4. The video shows the z axis of azoomed-in part of the three expanded cyst walls facing the host cell environment (left hand side). The arrows indicate the direction of movement along z axis. The “up” arrow indicate the movement away from the glass coverslip towards the top of the cyst, and the “down” arrow vice versa. The video demonstrates GRA2 puncta “spanning” in or around the CST1-positive layer of the cyst wall.

